# Transcriptional Regulator, Metabolic, and Pilus Biosynthesis Genes as Candidate Virulence Markers in High-Virulence *Mycobacterium abscessus*

**DOI:** 10.1101/2025.05.08.652910

**Authors:** Megan J. Maddox, Takehiro Kado

## Abstract

*Mycobacterium abscessus* is a rapidly growing non-tuberculosis mycobacterium that opportunistically causes pulmonary infections and is notable for its high resistance to antibiotics. While ubiquitous in environmental reservoirs such as soil, water, and biofilms, the genetic factors that enable certain strains to invade and persist in human hosts have remained elusive. To address this gap, we compared whole-genome sequences from 45 environmental isolates— primarily collected on Hawaiʻi Island—with a globally sourced set of clinical isolates retrieved from NCBI RefSeq. Our phylogenetic reconstruction delineated environmental and clinical lineages, and pangenome profiling revealed a conserved core genome of approximately 4,800 genes alongside a large accessory genome. Crucially, we found 20 genes conserved in clinical isolates, but not environmental isolates. Clinical isolates were found to have conserved genes coding for *erm(41)*, three transcriptional regulators, a pilus synthesis gene, an NADP-dependent oxidoreductase, a carbonic anhydrase, a probable L-ectoine synthase, a ribonuclease P protein component, and 11 hypothetical genes, whereas environmental isolated did not conserve any of these genes. This indicates that clinical isolates have undergone broad changes relating to metabolism, pilus biosynthesis, and transcriptional regulators, which may be necessary for pathogenicity. The candidate virulence markers uncovered here lay the groundwork for experimental validation, rapid diagnostics for non-tuberculous mycobacteria, and targeted surveillance of environmental reservoirs to mitigate the emergence of clinically significant strains.

**Importance:** *Mycobacterium abscessus* is an emerging pathogen that causes pulmonary infections in susceptible individuals. Though currently an opportunistic pathogen, *M. abscessus* may be undergoing evolutionary changes to become an obligate pathogen. Because of this, there is the need to identify markers of virulence in *M. abscessus* in order to better screen isolates and assess their risk of causing infection. This also provides the opportunity to study a similar pathogenesis to what *Mycobacterium tuberculosis* may have undergone. This study revealed that clinical isolates of *M. abscessus* have conserved genes relating to pilus biosynthesis, metabolism, and transcriptional regulators that environmental isolates do not have. These conserved genes may be required for *M. abscessus* to acquire virulence, and these genes have potential use as biomarkers to screen isolates for possible pathogenicity.

## Introduction

*Mycobacterium abscessus* is currently an emerging opportunistic pathogen that is known to cause pulmonary and soft tissue infections, and those with Cystic Fibrosis and surgical wounds are especially vulnerable (1). It is a rapidly growing non-tuberculosis mycobacterium that possesses a notably high resistance to antibiotics (2). The standard treatment for *M. abscessus* infection involves a 12 to 18 month antibiotic regimen but only results in a clearance rate of ∼45% (3). Although *M. abscessus* is ubiquitous in environmental reservoirs—such as soil, water distribution systems, and dusts (4)—the factors that enable environmental isolates to acquire high virulence and the evolutionary processes driving its emergence as a human pathogen remain poorly understood.

Of the isolates of *M. abscessus* found in clinical cases, 70% have been identified as belonging to 3 dominantly circulating clones, which are genetically clustered, highly virulent clones with greater rates of antibiotic resistance than non-clustered isolates (5). It was previously hypothesized that this genetic clustering of *Mycobacterium abscessus* was caused by patient-to-patient transmission, a claim that has been refuted by epidemiological studies (6). Due to there currently being little evidence for person-to-person transmission, it is generally accepted that infections with non-tuberculosis mycobacteria are acquired from environmental exposure to an opportunistic pathogen (7). It has been proposed that the genetic clustering seen in clinical isolates of *M. abscessus* is instead the result of mutations in the DNA repair machinery which slowed the rate of mutation (6). This opens up the possibility that clinical *M. abscessus* strains may have to acquire the genes necessary for pathogenicity while in the environment.

By leveraging the expanding genomic resources available in public databases, we compared the genomes of 45 environmental *M. abscessus* isolates—primarily from Hawaiʻi (4, 8)—with clinical isolates of *M. abscessus* collected worldwide (NCBI RefSeq). Our analysis identified 20 proteins uniquely conserved in clinical strains, most of which are involved in gene regulation and metabolic pathways. These results suggest that environmental isolates must acquire specific regulatory and metabolic functions to evolve into pathogenic forms capable of causing severe lung disease in cystic fibrosis patients. This study therefore aimed to pinpoint genes present in clinical—but absent from environmental—isolates in order to uncover the molecular determinants required for *M. abscessus* pathogenicity.

## Results

### Thirty-seven of the Forty-five Environmental Isolates Form a Clade with Dominantly Circulating Clones

The purpose of this study was to analyze the genomes of clinical and environmental isolates of *Mycobacterium abscessus* in order to find genes conserved in the clinical but not environmental isolates. This was done with the goal of finding genes which promote pathogenicity. For this study, 45 whole-genome sequences were used to compare clinical and environmental isolates (Table 1), all of which were obtained in previous research (1). The raw sequences were processed, assembled, and annotated, and these annotated genomes were then used for further analyses. The genomes of the 45 environmental isolates were put into KBase Insert Genome Into SpeciesTree v2.2.0 (9), which was then set to produce a phylogenetic tree with the isolates’ 200 closest relatives by analyzing 41 core house-keeping genes. This revealed that the isolates formed a clade with *M. abscessus* (Figure 1, left), confirming that the environmental isolates are *M. abscessus*. To obtain further information about these environmental isolates, a second phylogenetic tree was produced using the public genome information of clinical isolates of *M. abscessus* in NCBI RefSeq (10–12). There are certain *M. abscessus* strains of dominantly circulating clones (DCCs), which are responsible for over 70% of *M. abscessus* infections (5). DCCs are groups of genetically clustered isolates of *M. abscessus* that have been shown to display increased virulence in mice (5). By adding the known isolates DCC1–3 (ATCC19977, 3A-0119-R, 4S-0116-R, 6G-0125-R, 9808, 5S-0817, 1S-151-0930, 2B-0107, and 1049 (13)) to this phylogenetic tree, we found that 29 strains out of 36 formed a clade with DCC1 (Figure 1, right). Based on these data, the isolates classified with DCC strains may have been introduced to the environment after causing infection rather than being environmental strains of *M. abscessus*. Because of this, the study focused on the seven environmental isolates which were not part of DCC1.

**Figure 1:**
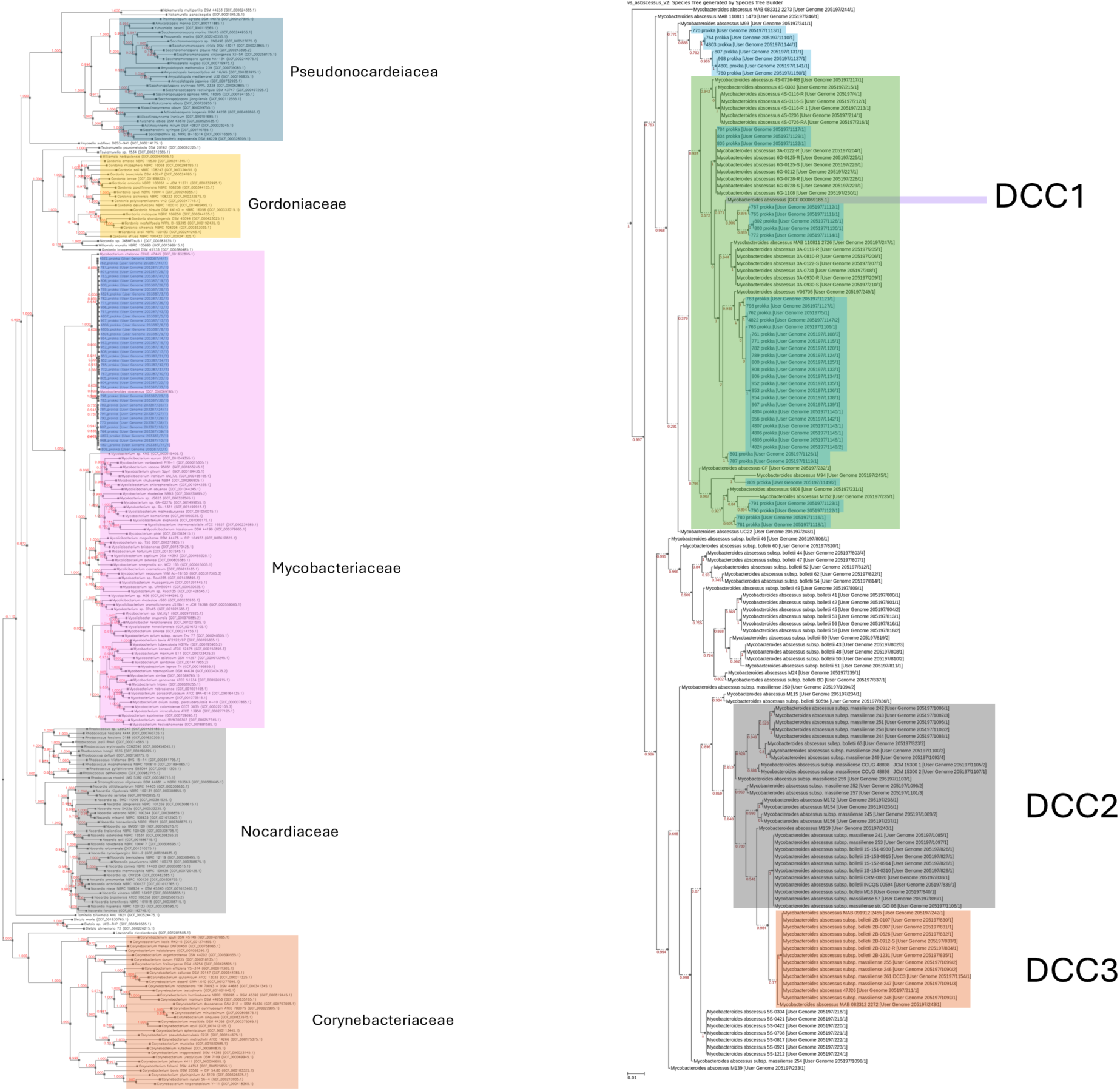
Phylogenetic relationships of *Mycobacterium abscessus* environmental isolates and 200 closest relative species. The phylogenetic tree was constructed based on 49 core genes using KBase Insert Genome Into SpeciesTree v2.2.0. Species in the family Pseudonocardeiacea are shadowed in blue, Gordoniaceae are shadowed in yellow, Mycobacteriaceae are shadowed in pink, Nocardiaceae are shadowed in gray, and Corynebacteriaceae were shadowed in orange. Environmental isolates which were analyzed are shadowed in blue. DCC1 lab strain ATCC19977 is shadowed in purple (left).

**Table 1:** Environmental isolates of *M. abscessus*.

| No. | Name | Accession | Library Name | Organism |
| --- | --- | --- | --- | --- |
| 1 | 760 | SRR15713760 | 12-48-SW-B-1.MAB | *Mycobacterium abscessus* subsp. *abscessus* |
| 2 | 761 | SRR15713761 | 12-42-SW-B-3.MAB | *Mycobacterium abscessus* subsp. *abscessus* |
| 3 | 762 | SRR15713762 | 12-39-Sw-B-1.MAB | *Mycobacterium abscessus* subsp. *abscessus* |
| 4 | 763 | SRR15713763 | 12-39-SW-A-1.MAB | *Mycobacterium abscessus* subsp. *abscessus* |
| 5 | 764 | SRR15713764 | 12-9-SW-B-2.MAB | *Mycobacterium abscessus* subsp. *abscessus* |
| 6 | 765 | SRR15713765 | RESP-ASH-D3-37 | *Mycobacterium abscessus* subsp. *abscessus* |
| 7 | 767 | SRR15713767 | RESP-ASH-D3-30 | *Mycobacterium abscessus* subsp. *abscessus* |
| 8 | 770 | SRR15713770 | KPH-43 | *Mycobacterium abscessus* subsp. *abscessus* |
| 9 | 771 | SRR15713771 | KPH-39 | *Mycobacterium abscessus* subsp. *abscessus* |
| 10 | 772 | SRR15713772 | FINE-ASH-D7-2-30 | *Mycobacterium abscessus* subsp. *abscessus* |
| 11 | 780 | SRR15713780 | 19-M-JP-19-F-1-37 | *Mycobacterium abscessus* subsp. *abscessus* |
| 12 | 781 | SRR15713781 | 19-M-JP-19-F-1-30 | *Mycobacterium abscessus* subsp. *abscessus* |
| 13 | 782 | SRR15713782 | 19-KPH-67-T-1-37 | *Mycobacterium abscessus* subsp. *abscessus* |
| 14 | 783 | SRR15713783 | 19-KPH-54-B-2-37 | *Mycobacterium abscessus* subsp. *abscessus* |
| 15 | 784 | SRR15713784 | 19-KPH-54-B-1-30 | *Mycobacterium abscessus* subsp. *abscessus* |
| 16 | 787 | SRR15713787 | 18-T-UHH-1-B-1-37 | *Mycobacterium abscessus* subsp. *abscessus* |
| 17 | 789 | SRR15713789 | 18-T-UHH-1-B-1-30 | *Mycobacterium abscessus* subsp. *abscessus* |
| 18 | 790 | SRR15713790 | 18-M-KF-6-W-1-37 | *Mycobacterium abscessus* subsp. *abscessus* |
| 19 | 791 | SRR15713791 | 18-M-KF-6-W-1-30 | *Mycobacterium abscessus* subsp. *abscessus* |
| 20 | 792 | SRR15713792 | 18-M-KF-6-W-1-22 | *Mycobacterium abscessus* subsp. *abscessus* |
| 21 | 798 | SRR15713798 | 17-17-SW-A-1-2-30 | *Mycobacterium abscessus* subsp. *abscessus* |
| 22 | 800 | SRR15713800 | 17-17-SW-A-1-1-30 | *Mycobacterium abscessus* subsp. *abscessus* |
| 23 | 801 | SRR15713801 | 17-137-SW-A-1-1-30 | *Mycobacterium abscessus* subsp. *abscessus* |
| 24 | 802 | SRR15713802 | 17-129-Sw-A1-1-37.MAB | *Mycobacterium abscessus* subsp. *abscessus* |
| 25 | 803 | SRR15713803 | 17-129-SW-A-1-1-30 | *Mycobacterium abscessus* subsp. *abscessus* |
| 26 | 804 | SRR15713804 | 17-11-SW-A-1-1-37 | *Mycobacterium abscessus* subsp. *abscessus* |
| 27 | 805 | SRR15713805 | 17-11-SW-A-1-1-30 | *Mycobacterium abscessus* subsp. *abscessus* |
| 28 | 806 | SRR15713806 | 12-9-SW-B-3.MAB | *Mycobacterium abscessus* subsp. *abscessus* |
| 29 | 807 | SRR15713807 | 12-57-SW-A-1.MAB | *Mycobacterium abscessus* subsp. *abscessus* |
| 30 | 808 | SRR15713808 | 12-50-SW-A-1.MAB | *Mycobacterium abscessus* subsp. *abscessus* |
| 31 | 809 | SRR15713809 | 12-48-SW-B-3 | *Mycobacterium abscessus* subsp. *abscessus* |
| 32 | 952 | SRR26031952 | 17N-17-Sw-B1-1-37.MAB | *Mycobacterium abscessus* subsp. *abscessus* |
| 33 | 953 | SRR26031953 | 17-51-Sw-B1-1-37.MAB | *Mycobacterium abscessus* subsp. *abscessus* |
| 34 | 954 | SRR26031954 | 17-17-Sw-A1-1-37.MAB | *Mycobacterium abscessus* subsp. *abscessus* |
| 35 | 956 | SRR26031956 | 17-15-Sw-A1-1-37.MAB | *Mycobacterium abscessus* subsp. *abscessus* |
| 36 | 967 | SRR26031967 | 12-45-Sw-A-2.MAB | *Mycobacterium abscessus* subsp. *abscessus* |
| 37 | 968 | SRR26031968 | 12-39-SW-B-2.MAB | *Mycobacterium abscessus* subsp. *abscessus* |
| 38 | 4801 | SRR25404801 | 12-39-SW-B-2.MAB | *Mycobacterium abscessus* |
| 39 | 4803 | SRR25404803 | 12-9-SW-B-2.MAB | *Mycobacterium abscessus* |
| 40 | 4804 | SRR25404804 | 17-15-Sw-A1-1-37.MAB | *Mycobacterium abscessus* |
| 41 | 4805 | SRR25404805 | 17-17-Sw-A1-1-37.MAB | *Mycobacterium abscessus* |
| 42 | 4806 | SRR25404806 | 17-51-Sw-B1-1-37.MAB | *Mycobacterium abscessus* |
| 43 | 4807 | SRR25404807 | 17N-17-Sw-B1-1-37.MAB | *Mycobacterium abscessus* |
| 44 | 4822 | SRR25404822 | 12-39-Sw-B-1.MAB | *Mycobacterium abscessus* |
| 45 | 4824 | SRR25404824 | 12-45-Sw-A-2.MAB | *Mycobacterium abscessus* |

### Core Genome Has Moderately Low Conservation Between Environmental and Clinical Isolates

Seven environmental isolates were obtained, and a phylogenetic tree was constructed using these isolates and publicly available clinical isolates of *M. abscessus*. To further investigate gene co-occurrence, we performed a pangenome-based phylogenetic analysis. This analysis included *Mycobacterium tuberculosis* H37Rv, three clinical isolates of *M. abscessus* (DCC 1–3), and the seven environmental isolates (Figure 2). Node 0 compared the genomes of *M. tuberculosis* and all isolates of *M. abscesses* and showed that, of the 10,024 genes analyzed, 23.4% were classified as singleton (present in only one genome), 56.2% as partial (present in more than one but fewer than all genomes), and 20.4% as core (present in all genomes) (Figure 2). This indicated a low level of conservation of the core genome between *M. tuberculosis* and *M. abscesuss*. Node 6 compared genomes of clinical isolates and the 7 environmental isolates of *M. abscessus* and showed that, of the 7770 genes analyzed, 24.7% were classified as singleton, 25.0% as partial, and 50.3% as core (Figure 2). Relative to nodes 3 (75.5% core) and 7 (71.0% core) (Figure 2), which compared isolates in DCC1 and DCC2 respectively, the core genome of *M. abscessus* had a moderately low level of conservation between DCC1 and DCC2, as well as between the environmental and clinical isolates.

**Figure 2:**
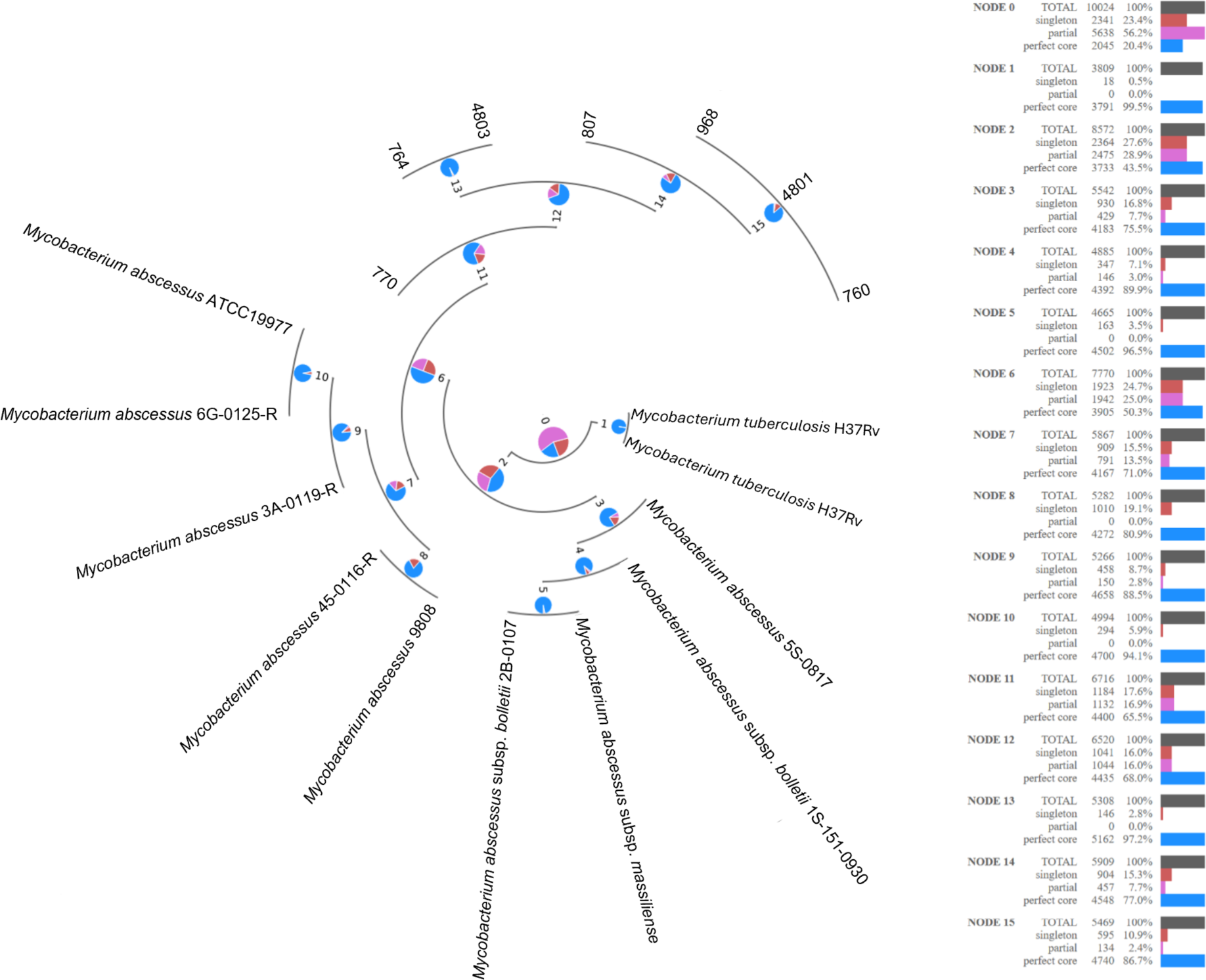
Pangenome-based phylogenomic analysis of *Mycobacterium abcessus* isolates and *Mycobacterium tuberculosis* H37Rv. Orthologous gene sets within the pangenome are divided into three categories: Core is represented by blue, Singleton by red, and Partial by pink. Pangenome-based phylogenomic analysis was created using KBase Phylogenetic Pangeonme Accumulation – v.1.4.0.

These data were influenced by the lower quality of next-generation sequencing. The assembled genes were assessed using QUAST (Quality Assessment Tool for Genome Assemblies (14, 15)). The results showed that isolates 770, 764, and 4803 had more than 500 contigs, while isolates 807 and 760 had over 180 contigs (Supplementary figure). In contrast, isolates 968 and 4801 had only 27 contigs (Supplementary figure), which is a reasonable number for a GC-rich organism like *Mycobacterium*. Given this, we decided to further analyze ortholog conservation between the environmental isolates (968 and 4801) and the highly virulent DCC strains.

### Conserved Genes Influence Regulators, Pilus Biosynthesis, and Metabolism

To identify genes essential for the pathogenicity of *M. abscessus*, we analyzed gene conservation between environmental isolates and highly virulent DCC strains. All genes were subjected to OrthoMCL, a genome-scale algorithm that clusters orthologous protein sequences, which are defined as those sharing a common ancestor. As a result, 21 genes were identified as being conserved in the clinical isolates of *Mycobacterium abscessus* that were not conserved in either environmental isolate (Table 2). Since *erm*(41), a gene strongly associated with high antimicrobial resistance in *M. abscessus* clinical isolates (16), is absent in both environmental isolates, its acquisition may be necessary for environmental isolates to become pathogenic. In addition to *erm*(41), three transcriptional regulators (3, 17, 18), a pilus synthesis gene (19), an NADP-dependent oxidoreductase (20), a carbonic anhydrase (21), a probable L-ectoine synthase (22), a ribonuclease P protein component (23), and 11 hypothetical genes were identified. These genes not being conserved in environmental isolates indicated that clinical isolates have undergone broad changes regarding regulators, pilus biosynthesis, and metabolism, which may be required for infection and survival within the host (24).

**Table 2:** Genes conserved in clinical isolates of *Mycobacterium abscessus* and not environmental isolates along with their descriptions.

| Gene Locus | Gene Description | Ref. |
| --- | --- | --- |
| MAB_2297 | 23S rRNA (adenine(2058)-N(6))-methyltransferase Erm(41) | (16) |
| MAB_1873c | Putative TetR family transcriptional regulator | (18) |
| MAB_1430c | Putative transcriptional regulator, TetR family | (18) |
| MAB_3446 | Putative transcriptional regulator, WhiB family | (3, 17) |
| MAB_0474 | pilus biosynthesis protein TadE | (19) |
| MAB_1431 | Conserved hypothetical protein (Putative monooxygenase) | (26) |
| MAB_1874 | NADP-dependent oxidoreductase | (20) |
| MAB_0559 | Probable carbonic anhydrase (carbonate dehydratase) | (21) |
| MAB_2585 | Probable L-ectoine synthase | (22) |
| MAB_4954c | ribonuclease P protein component | (23) |
| MAB_4666c | DUF1326 domain-containing protein | - |
| MAB_3411 | DUF2200 domain-containing protein | - |
| MAB_0093 | DUF3298 domain-containing protein | - |
| MAB_4592c | DUF456 domain-containing protein | - |
| MAB_0012c | hypothetical protein | - |
| MAB_1301c | hypothetical protein | - |
| MAB_1657c | hypothetical protein | - |
| MAB_2500 | hypothetical protein | - |
| MAB_3264c | hypothetical protein | - |
| MAB_4291 | hypothetical protein | - |
| MAB_4593c | hypothetical protein | - |

Among the genes conserved in the clinical isolates was one coding for a carbonic anhydrase, a type of enzyme that forms bicarbonate by catalyzing the hydration of carbon dioxide (21). This reaction is necessary for several processes, such as fatty acid biosynthesis and the production of certain small molecules (21), indicating differences in the metabolism of the clinical isolates due to the presence of an enzyme of this class. An L-ectoine synthase was also among the genes not conserved in the environmental isolates. In *Mycobacterium smegmatis*, ectoine synthesis has been shown to protect against slowed growth in saline environments (22), though currently its function has not been studied in *M. abscessus*. Lastly, a gene coding for the pilus biosynthesis protein TadE was found to be conserved in the clinical isolates of *M. abscessus*. In *Mycobacterium tuberculosis*, *tadE* codes for minor pilins, and its expression, along with all *tad/flp* genes, is up-regulated upon infection of macrophages and pneumocytes (19). This indicated that TadE may be required for *M. abscessus* to establish infection within host cells.

Erm(41) was also among these conserved genes and codes for a macrolide-inducible ribosomal methylase, which is responsible for macrolide resistance in *M. abscesses* (16). A WhiB family transcriptional regulator was also found among these conserved genes. *whiB7* is a member of this family of regulators and is the activator of *erm(41)* (17). Along with their activation of *erm*(*41*), MmpS-Mmpl efflux pumps dependent upon these regulators may contribute to the antibiotic resistance that is associated with this family of transcriptional regulators (17, 18). Previous research has shown that in *Mycobacterium bovis* BCG, the TetR family transcriptional regulator BCG0878c interacts with 3-methyladenine DNA glycosylase to promote genomic stability, assisting in the slowed mutation rate that caused the genetic clustering of clinical *Mycobacterium abscessus* (6, 25), though further studies are needed to confirm if a similar mechanism exists in *M. abscessus*.

NADP-dependent oxidoreductases and monooxygenases are large classes of enzymes with a wide range of functions in bacterial cells (20, 26). Mycobacteria require the substantial use of enzymes in the oxidative processes involved in ATP generation due to being obligate aerobic bacteria (27). Because of this, previous research has predicted that mycobacteria would have ∼200 oxidoreductases, with many of these oxidoreductases being involved ATP hydrolysis and the electron transport system (27).

### Key Transcription Factors Are Absent in Non-Pathogenic M. abscessus

To predict how the 21 genes identified by OrthMCL impact virulence of *M. abscessus*, these genes were searched in Mycobrowser to obtain their amino acid sequence. This amino acid sequence was then entered into STRING to find predicted protein-protein interactions (Figure 3). In addition, the orthologues of these genes within *Mycobacterium tuberculosis* H37Rv were searched in the Mtb transposon sequencing database (MtbTnDB), a central repository of TnSeq screens performed in *M. tuberculosis* to find information relating to the function of those genes in the pathogenicity of *M. tuberculosis* H37Rv. The orthologs MAB_1873, MAB_1430, MAB_0474, and MAB_4291 showed negative Log₂ fold change (Log₂FC) values and q-values below 0.05 in multiple Tn-seq analyses. According to database information, the environmental isolates lacking these four genes also lack isoniazid resistance and the ability to proliferate in the host (Table 3). Overall, our comparative gene analysis between environmental and clinical isolates led to the identification of environmental strains that do not recur in patients, as well as key transcriptional regulators missing in non-pathogenic *M. abscessus*. The acquisition of these factors, either from the environment or under complex infection conditions, may enable environmental *M. abscessus* strains to cause highly virulent, contagious lung disease.

**Figure 3:**
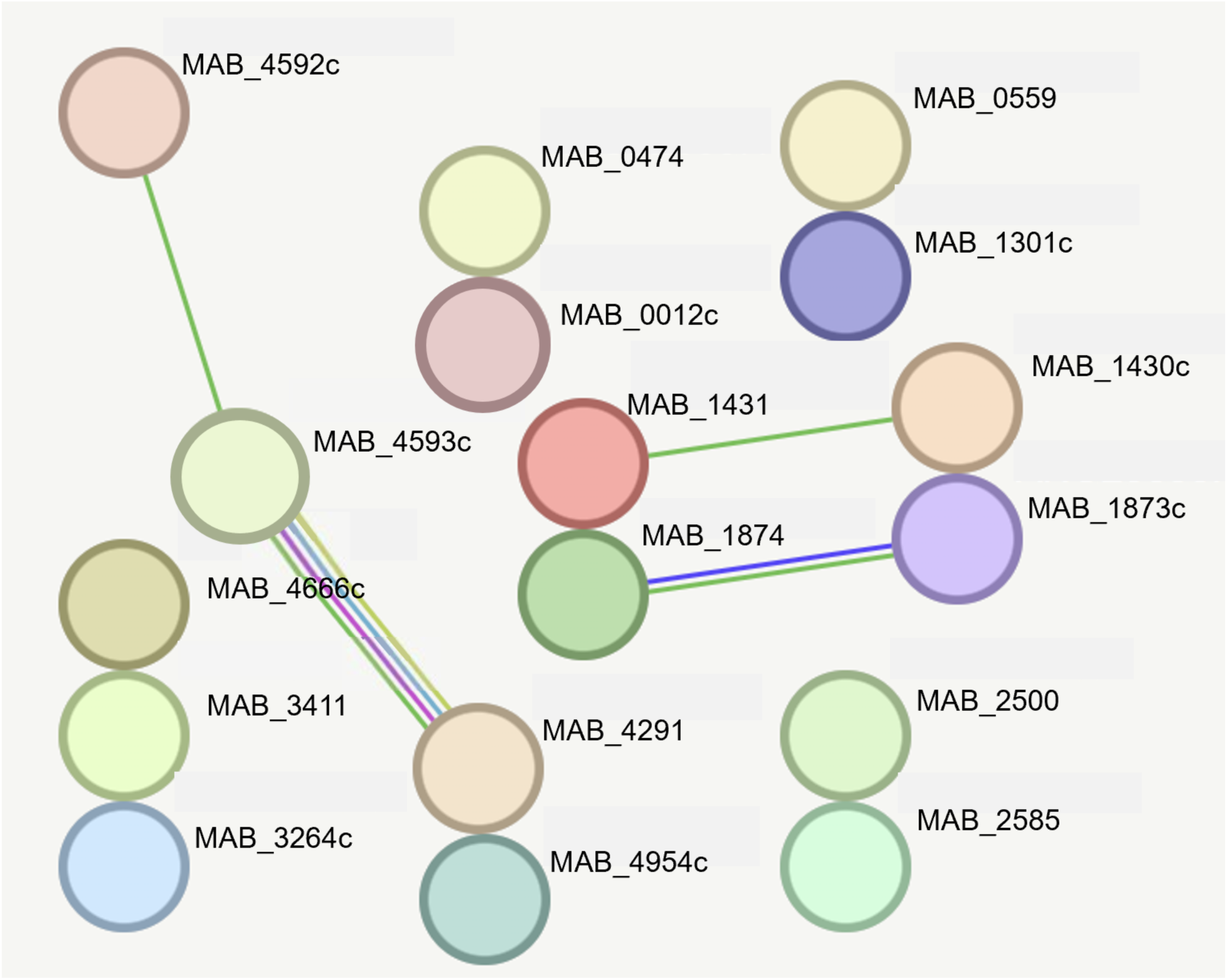
Protein-protein interactions between the analyzed genes as predicted by STRING. The nature of interactions is indicated by edge colors: known interactions are represented by blue (from curated databases) and purple (experimentally determined); predicted interactions are represented by green (gene neighborhood), orange (gene fusions), and dark blue (gene co-occurrence); other interactions are represented by olive (textmining), black (co-expression), and lilac (protein homology). Empty nodes are proteins with an unknown 3D structure, and filled nodes are proteins where the 3D structure is known or predicted.

**Table 3:** Data obtained from MtbTnDB regarding orthologues of the analyzed *Mycobacterium abscessus* genes in *Mycobacterium tuberculosis*.

| MAB Gene Locus | Orthologue in *M. tuberculosis* | Log2FC | q-val | *in vitro/ in vivo* | Meaning | Microarray or TnSeq | Ref. |
| --- | --- | --- | --- | --- | --- | --- | --- |
| MAB_1873c | Rv0067c | -1.65 | 0.04 | *in vitro* | Mutants exhibiting altered fitness in presence of isoniazid | TnSeq | (39) |
| MAB_0474 | Rv3655c | -1.39 | 0.02 | *In vivo* | Differential genetic requirement of Mtb (H37Rv) in mouse relative to *in vitro* | TnSeq | (40) |
| MAB_4291 | Rv2264c | -0.7 | 0 | *In vivo* | Differential genetic requirement of Mtb (H37Rv) in mouse relative to *in vitro* | TnSeq | (40) |

## Discussion

The increasing incidence of *Mycobacterium abscessus* infections in both cystic fibrosis and non–cystic fibrosis patients (1) underscores a critical gap in our understanding of how environmental populations give rise to clinically relevant strains. To address this, we performed deep whole-genome sequencing of environmental isolates alongside publicly available clinical genomes, reconstructed their evolutionary relationships using KBase’s *Insert Genome into SpeciesTree*, and conducted a comprehensive pangenome analysis with OrthoMCL and Phylogenetic Pangenome Accumulation. Our results demonstrate that environmental isolates cluster into distinct, well-supported clades separate from clinical lineages. Clinical lineages of *Mycobacterium abscessus* possess a large accessory genome enriched for 20 conserved genes not found in environmental isolates. Those genes included three genes involved in antibiotic resistance, four genes involved in diverse metabolic functions, and one involved in pilus biosynthesis. By distinguishing core from accessory gene repertoires, we show that environmental *M. abscessus* isolates carry genetic elements that may serve as reservoirs for future virulence or resistance traits. This insight into the evolutionary trajectory from environmental niche to human pathogen also pinpoints 20 candidate molecular markers of high virulence in *M. abscessus*.

One restriction in this study was the limited analyses that could be conducted on the environmental isolates of *Mycobacterium abscessus* studied in this research. Even among clinical isolates of mycobacteria, equivalent virulence cannot be assumed due to the intraspecies differences of isolates (1). Previous research showed that non-pathogenic mycobacteria induced a greater TNFα response in human macrophages than pathogenic mycobacteria and found that *M. abscessus* isolates which produced a rough colony morphology were more infections than those with a smooth morphology (28). Another previous study found two environmental *M. abscessus* isolates displayed comparable survival to a clinical isolate in THP-1 macrophages (29). However, these studies did not include isolate 798 (Fig. 2). Because of this and the broad intraspecies differences in mycobacterial virulence, it is currently unknown whether the environmental isolates of *M. abscessus* used in this study truly exhibit less virulence than the clinical isolates or how certain conserved proteins in the clinical isolates contribute to their pathogenicity. Future experimental validation will be essential to resolve these questions.

Genes encoding Erm(41) and transcriptional regulators from the WhiB and TetR families contribute to pathogenesis and antibiotic resistance in *M. abscessus* (3, 16–18). Our identification of these regulators as hallmarks of highly virulent strains echoes previous reports of significant enrichment in genes associated with transcriptional control and DNA modification within dominant circulating clones (DCCs) versus non-DCC isolates (13). Horizontal gene transfer provides a plausible route for the introduction of these resistance and regulatory elements into *M. abscessus*, potentially driving the rapid emergence of antibiotic-resistant, virulent lineages. Horizontal gene transfer has been implicated in mycobacteria obtaining traits such as antibiotic resistance and pathogenesis (27). A previous study found that in horizontal gene transfer to *Mycobacteria*, potential donor organisms were mainly soil-inhabiting organisms, indicating that the environment can be an important site for the transfer of genetic material (27). This same study analyzing horizontally transferred genes (HGTs) in mycobacterium found that genes relating to metabolic pathways and enzymatic activity were over-represented among candidate HGTs (27). This is consistent with the results of our study. Because the genes found to be conserved in clinical *M. abscessus* largely relate to enzymatic functions, this suggests that these genes were potentially acquired by horizontal gene transfer in the environment.

The functions of particular proteins, such as TadE and L-ectoine, have currently not been studied specifically within *M. abscessus*. Furthermore, certain classes of enzymes found to be conserved in clinical isolates have wide ranges of potential functions, and the specific enzyme conserved could not be determined. This is indicative of the need for further experimentation to investigate the functions of these proteins and enzymes in *M. abscessus* and to ascertain the specific differences between the clinical and environmental isolates that contribute to pathogenicity.

In summary, the conserved transcriptional regulators we identified in clinical *M. abscessus* isolates—including TetR- and WhiB-family proteins—alongside TadE, L-ectoine biosynthesis enzymes, and Erm(41), are all intimately linked to antibiotic resistance and may be essential for environmental strains to acquire pathogenic potential. Because rapid assays for non-tuberculous mycobacterial virulence are lacking, these genes represent promising biomarkers for screening environmental isolates and assessing their risk of causing disease. Validating these candidate virulence determinants in future experimental studies will be critical for the development of diagnostic tools and for informing public-health strategies to combat *M. abscessus* infections.

## Material and Method

### Processing of Raw Whole-Genome Sequence Data

All analyses and data were documented in narratives hosted on KBase (https://narrative.kbase.us/narrative/199945). Raw sequence data for environmental *Mycobacterium abscessus* isolates were downloaded from the Sequence Read Archive (SRA) at NCBI, as provided by Honda et al. (30). Quality control of raw reads from whole-genome sequencing was performed using FastQC v0.11.7 (31). Subsequently, TruSeq Illumina adapters and low-quality, short sequences were removed using Trimmomatic v0.36 (32) and PRINSEQ v20. 4 (33). Additional read quality filtering was completed using Filtlong v0.2.1 (34). Genome assembly into contigs was carried out with SPAdes v3.15. 3 (35, 36), and the assembled genomes were annotated using Prokka v1.14.5 (37).

### Phylogenetic Analysis

A phylogenetic tree incorporating 200 closely related species was constructed using KBase’s *Insert Genome into SpeciesTree* module (v2.2.0) (9), with the analysis archived in a KBase narrative (https://narrative.kbase.us/narrative/203387). For a comparative phylogenetic analysis of clinical *M. abscessus* isolates, annotated genomes were retrieved from the NCBI RefSeq database. The sequence data for both clinical and environmental isolates were processed using the same *Insert Genome into SpeciesTree* module (v2.2.0) (9), and the resulting phylogenetic tree was saved in another KBase narrative (https://narrative.kbase.us/narrative/205197).

### Metagenomic Analysis

All metagenomic analyses and corresponding data were managed within a KBase narrative (https://narrative.kbase.us/narrative/205969). A genome set for pangenome analysis was generated using the KBase *Insert Genome into SpeciesTree* module (v2.2.0) (9). The pangenome was constructed with OrthoMCL v2.0 (38), and core genome analysis was conducted using Phylogenetic Pangenome Accumulation v1.4.0 (38).

## Acknowledgement

This research received no specific grant from any funding agency in the public, commercial, or not-for-profit sectors. The authors thank OpenAI’s ChatGPT (GPT-4.0) for assistance with English-language polishing of the manuscript.

## Supplementary figure

**Supplementary figure.**
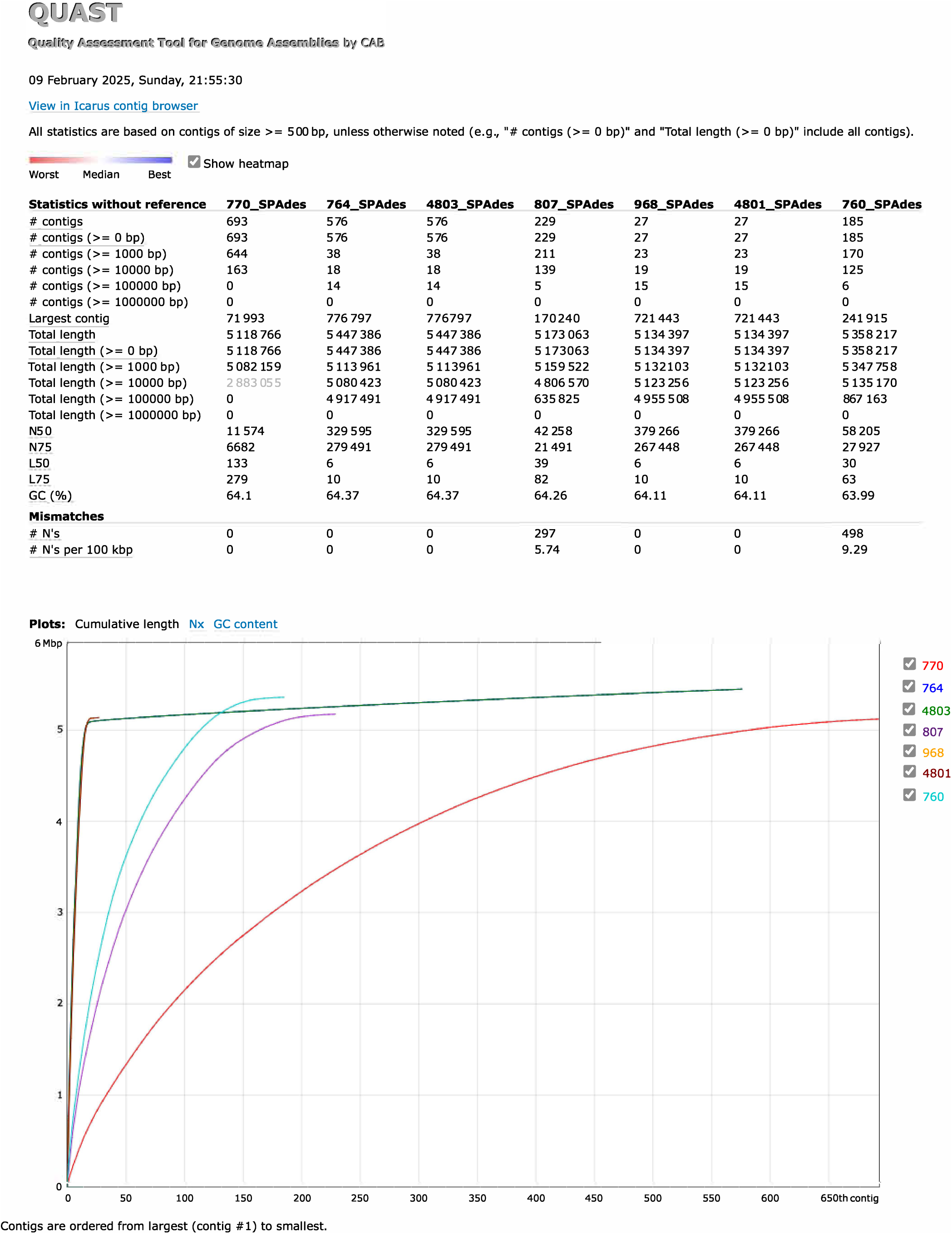
The only isolates 968 and 4801 had reasonable contigs. The assembled genome sequences of prospective environmental isolates of *M. abscessus* were subjected to the quality check by QUAST

